# Cell-wall synthesis and ribosome maturation are co-regulated by an RNA switch in *Mycobacterium tuberculosis*

**DOI:** 10.1101/232314

**Authors:** Stefan Schwenk, Alexandra Moores, Irene Nobeli, Timothy D. McHugh, Kristine B. Arnvig

## Abstract

The success of *Mycobacterium tuberculosis* as a pathogen relies on the ability to switch between active growth and non-replicating persistence, associated with latent TB infection. Resuscitation promoting factors (Rpfs) are essential for the transition of *M. tuberculosis* to dormancy and for emergence from the non-replicating persistent state. But these enzymes are double-edged swords, as their ability to degrade the cell wall, is potentially lethal to the bacterium itself. Hence, Rpf expression is tightly regulated. We have identified a novel regulatory element in the 5’ untranslated region (UTR) of *rpfB*. We demonstrate that this element is a transcriptionally regulated RNA switch/riboswitch candidate, which is restricted to pathogenic mycobacteria, suggesting a role in virulence. Moreover, we have used translation start site mapping to re-annotate the RpfB start codon and identified and validated a ribosome binding site that is likely to be targeted by an RpfB antisense RNA. Finally, we show that *rpfB* is co-transcribed with downstream genes, *ksgA* and *ispE. ksgA* encodes a universally conserved methyl transferase involved in ribosome maturation and *ispE* encodes an essential ATP-dependent kinase involved in cell wall synthesis. This arrangement implies co-regulation of resuscitation, cell wall synthesis and ribosome maturation via the RNA switch. We propose that deregulation of this switch, associated with cell wall synthesis and ribosome function, presents a new target for anti-tuberculosis drug development.

**Importance:** This work describes the identification and characterisation of a novel regulatory RNA element/attenuator that controls cell wall synthesis and ribosome function in *Mycobacterium tuberculosis*, the causative agent of human tuberculosis (TB). By switching between two different conformations, this RNA switch can either enable or inhibit transcription of a tri-cistronic mRNA that encodes a cell-wall remodelling enzyme crucial for activation of latent TB, an RNA methytransferase that is important for ribosome function and a protein kinase essential for early steps in cell wall synthesis. This RNA switch is only present in a subset of pathogenic mycobacteria, and by regulating the expression of three genes associated with classical antimicrobial targets we believe that it offers a novel important target for future anti-tuberculosis drugs.

## Introduction

The ability to switch between actively replicating and non-replicating persistence (NRP) is at the heart of *Mycobacterium tuberculosis’* success as a pathogen. *M. tuberculosis* expresses five resuscitation-promoting factors (RpfA-E) (1). These are cell wall remodelling enzymes critical for the transition of *M. tuberculosis* between dormancy and resuscitation, and for reactivation of tuberculosis (TB) in animal models (2-4). In an *in vivo* environment, *M. tuberculosis* forms cells that can only be grown with Rpf supplementation (5).

Precise and tight control of Rpf expression is vital as these enzymes are able to degrade the bacterial cell wall posing a potentially lethal threat to *M. tuberculosis* itself. Expression of the five Rpfs is induced by different triggers, many of which are associated with the host environment (6,7). ChIP-seq data indicates that several transcription factors, including MtrA and Lsr2 regulate these promoters (8).

RNA-based regulation (riboregulation) of bacterial gene expression has attracted increasing attention over the last decade, as the abundance of the molecules and the systems they regulate become increasingly obvious (9-14). One class of riboregulators are the RNA-switches, cis-regulatory elements, located largely within the 5’ untranslated region (UTR) of the mRNA they regulate. Upon sensing a physiological signal such as temperature, pH, metabolites, RNA or proteins, they switch between conformations that are either permissive or non-permissive for downstream gene expression; RNA switches regulated by small molecule ligands are specifically referred to as riboswitches, and these currently make up the largest class of RNA switches (13,15,16). Riboswitches are formed of distinct domains with an aptamer domain responsible for binding a specific ligand, and an expression platform that regulates transcription or translation downstream (17). Most of the riboswitches described to date are widespread and associated with biosynthetic pathways; however, there are examples of less widespread riboswitches, and it is likely that there are many more, some of which may never be identified due to their rare occurrence (17,18). Riboswitches have been highlighted as potential drug targets due to their inherent ability to interact with a variety of ligands. For example, the FMN riboswitch has been suggested as potential drug target against *M. tuberculosis* infection (19).

Here we identify a novel transcriptional RNA switch (riboswitch candidate) located within the 5’ UTR of *M. tuberculosis rpfB*, and restricted to a subset of pathogenic mycobacteria. Based on experimental evidence, we have re-annotated the RpfB start codon and identified a likely Shine-Dalgarno (SD) sequence (20) that overlaps with an asRNA transcribed opposite to RpfB. The genetic arrangement of *rpfB* flanked upstream by the *tatD* nuclease, and downstream by the universally conserved *ksgA* methyl transferase and the essential *ispE* kinase is conserved in a wide range of Actinobacteria (Fig. S1) (21). We show that *rpfB, ksgA* and *ispE* are co-transcribed indicating a tight regulatory link between resuscitation, cell wall synthesis and ribosome maturation, subject to regulation by this novel element.

## Results

### Promoters and transcripts of the *rpfB* locus

Through interrogation of *M. tuberculosis* (d)RNA-seq (22), we found that *rpfB* is expressed from two promoters: P1, with transcription start site (TSS) at G1127876, and P2 with TSS at A1127955. For both TSS we identified canonical (TANNNT) -10 regions (Fig. 1). RNA-seq also indicates the presence of an antisense RNA expressed from P_as_ with TSS at G1128048. Expression from these promoters was validated by cloning the region from 140 basepairs upstream of P1 to the annotated ATG start codon in frame to a *lacZ* reporter (Fig. 1). In addition, we made three derivatives mutating the -10 regions of either P1 or P2 separately or in both P1 and P2. Finally, we made a transcriptional fusion of P_as_ including 100 basepairs of upstream region. The constructs were transformed into *M. tuberculosis* and promoter activity was assessed by colony colour on X-gal plates (Fig. 1C). Mutating the promoters individually suggested that P1 and P2 are both active in *M. tuberculosis*, corroborating the RNA-seq data. The lack of expression in the double mutant supports the TSS mapping indicating that P1 and P2 are the only promoters driving *rpfB* expression. Moreover, the results indicate that Pas is active and likely to play a role in *rpfB* expression.

**Fig. 1:**
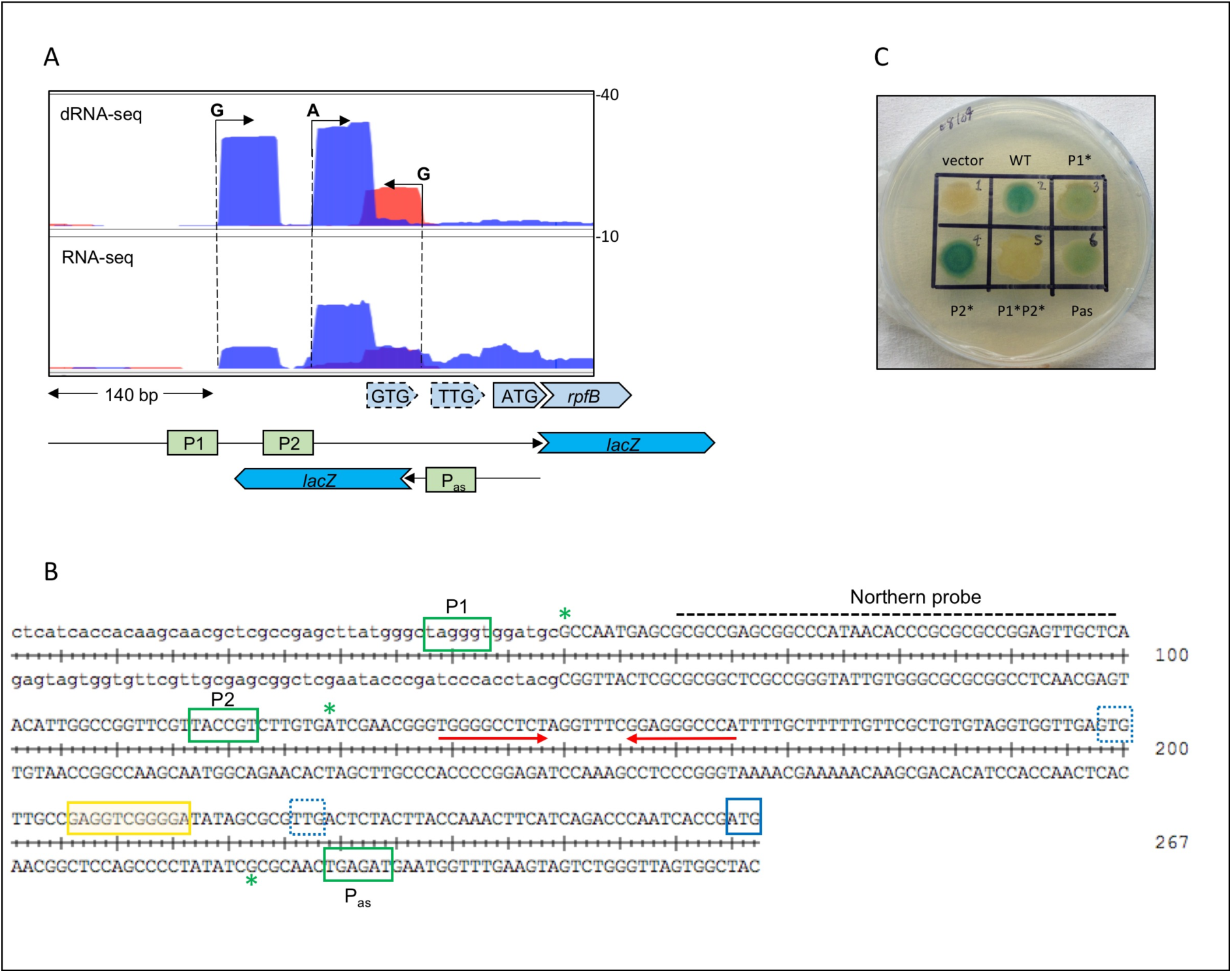
Transcription start sites (TSS) and promoter elements in the *rpfB* locus. A: dRNA-seq and RNA-seq of the promoter region, 5’ UTR and early ORF of *rpfB*. Numbers on the right indicate normalised reads. Three TSS were identified; two sense and one antisense (data from (22)). Below are schematics of the regions covered by the reporter constructs. B: Sequence of promoter region and 5’ UTR of *rpfB*. Green boxes indicate the -10 hexamers of promoters P1, P2 and P_as_; green asterisks: TSS. Red arrows: inverted repeat leading to a stem-loop structure; blue box: annotated translation start site, blue dotted boxes: alternative translation start sites, yellow box: putative ribosome binding site. C: X-gal plate with reporter constructs expressed in *M. tuberculosis*. Wildtype refers to 'full-length' construct from -140 upstream of P1 to the ATG. Mutations in the -10 region (TANNNT to CANNNC) are indicated by asterisk (i.e. P1* has inactivated P1, but functional P2 and *vice versa*).

### RpfB translation start site

The annotated translation start site of RpfB is ATG (Fig. 1). However, there is no obvious SD sequence proximal to this start codon; moreover, the start sites of the RpfB homologues in *Mycobacterium leprae* and *Mycobacterium smegmatis* have been annotated 13 codons further upstream, corresponding to the alternative TTG start (Fig. 1), which has a likely SD sequence upstream. In line with previous observation, we considered that the RpfB start codon may have been mis-annotated (23,24). We found two potential start sites (TTG and GTG) upstream of the annotated ATG (Fig. 1). In order to define which of the potential start sites was correct, we modified the method developed by Smollett et al. for translation start site mapping (24), using the wildtype translational *lacZ* fusion described above. Frameshift mutations were introduced separately between GTG and TTG and between TTG and ATG. If a frameshift were located within the resulting coding sequence, functional beta-galactosidase (β-gal) would not be expressed. The constructs were transformed into *M. smegmatis*, a tractable surrogate host for the expression of *M. tuberculosis* genes, and cell extracts were assayed for β-gal activity.

The results, shown in Fig. 2, demonstrate that the frameshift between GTG and TTG retained ~75% of wildtype β-gal activity level, suggesting that this part of the transcript was outside the translated region. However, the frameshift between TTG and ATG reduced β-gal activity to the level of the empty vector, indicating the mutation lay within the translated region and hence that TTG was the correct start codon (Fig. 2). As this result was in conflict with previously published data (25), we employed an alternative method to validate our findings. Each of the three potential start sites (GTG, TTG, ATG) was mutated to non-start codons (GTC, TTA, AAG), and β-gal activity of the resulting constructs assayed. The results (Fig. 2) corroborated our findings from the frameshift experiment; changing GTG and ATG to non-start codons did not significantly reduce β-gal activity, while changing the TTG to TTA reduced the expression to empty vector level, thus verifying that TTG was the correct start codon. Further supporting this notion was the fact that we could only identify a putative SD sequence -10 to -20 relative to TTG (Fig. 1B). To investigate if this sequence affected *rpfB* expression, we mutated the SD purines to pyrimidines in the *lacZ* fusion. The β-gal activity of the resulting construct was reduced to the level of the empty vector (SD mut, Fig. 2), suggesting that this was a likely ribosome binding site.

**Fig. 2:**
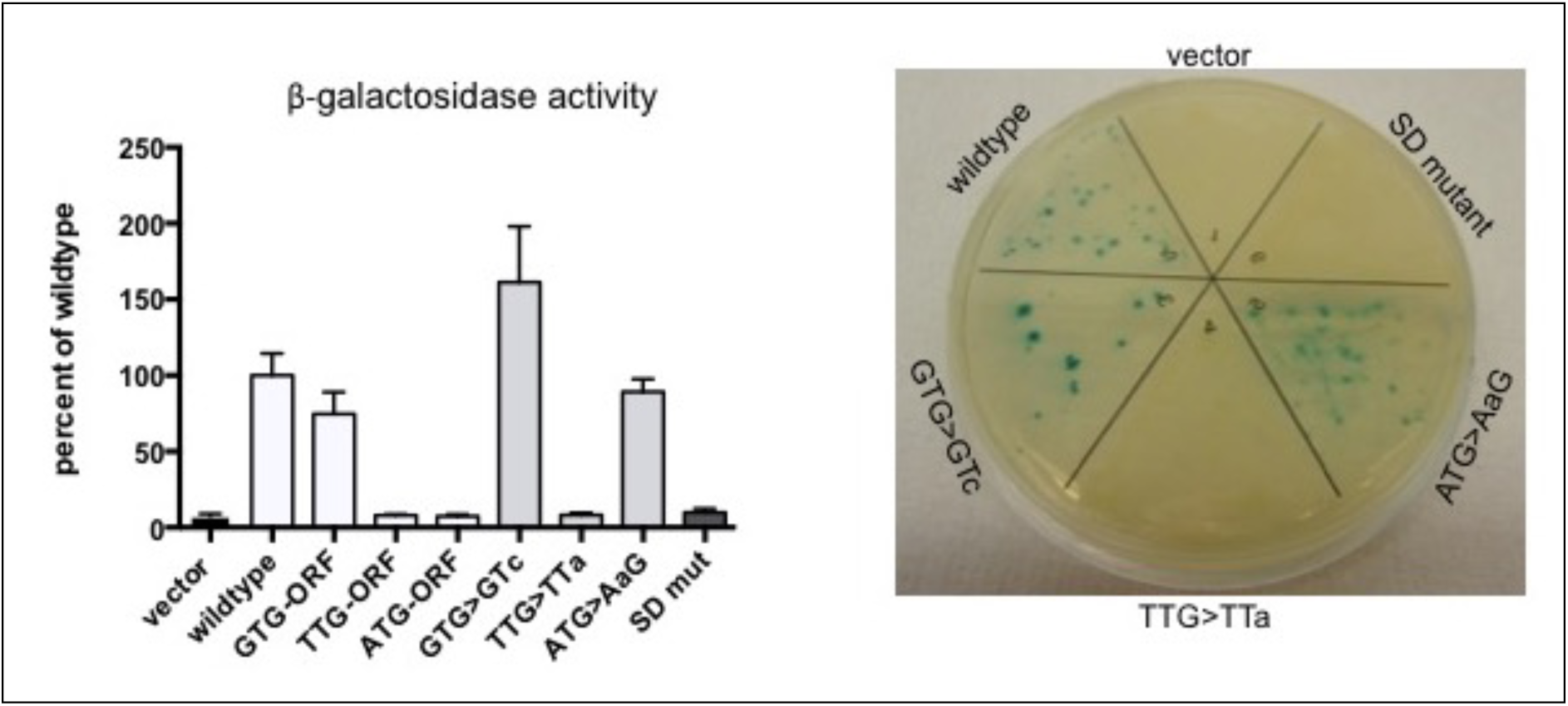
Translation start site mapping. Left panel shows results of β-gal assays on translational reporters expressed in *Mycobacterium smegmatis*. GTG-ORF, TTG-ORF, ATG-ORF indicate the results of introducing frameshifts in the open reading frames downstream of the indicated putative start site. GTG>GTc, TTG>TTa, ATG>AaG indicate the results of changing the putative start site to a non-start codon. The last bar in the graph shows the activity of the mutated SD sequence (GAGGTCGGGGA to ctccTCcccct). Right panel shows translation start site and SD mutants expressed in *M. tuberculosis*.

Finally, we transformed selected constructs with altered start sites into *M. tuberculosis* to ensure there were no significant differences compared to *M. smegmatis.* The results in *M. tuberculosis,* seen as blue/white colony colour (Fig. 2), were in perfect agreement with the results obtained in *M. smegmatis*, supporting the notion that the correct translation start site for *M. tuberculosis* RpfB is the relatively unusual TTG codon.

### The RpfB 5’ UTR

The first 130 nucleotides of the RpfB 5’ UTR expressed from P1 include an inverted repeat (red arrows, Fig. 1) followed by a poly-U tract, suggestive of a potential intrinsic terminator. Using *mfold* (26), we found that the predicted structure of the 130 nucleotides does indeed contain a stem-loop followed by a poly-U tract (Fig 3A). In order to determine if this sequence might lead to premature transcription termination, we analysed RNA from exponential and stationary phase cultures of *M. tuberculosis* and the closely related *Mycobacterium bovis* BCG by Northern blotting. Figure 3B shows a Northern blot with a strong signal around 125 nucleotides in exponential phase from both species, consistent with a terminated transcript. In addition, there are several weaker signals corresponding to larger transcripts. In stationary phase, there was little or no expression in both species, in concordance with previous observations (6,27).

**Fig. 3:**
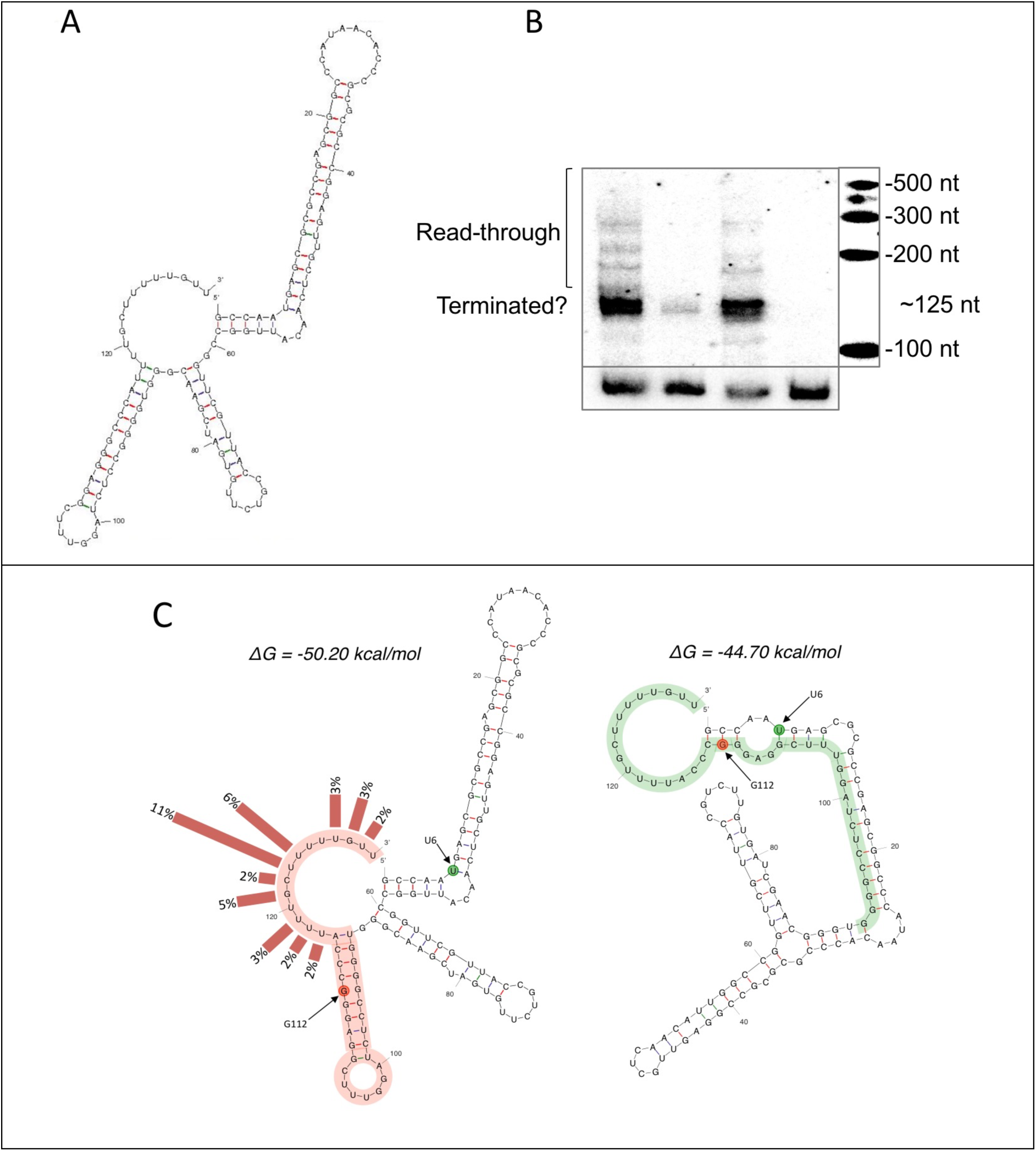
*rpfB* 5’ UTR. A: *mfold* (26) predicted structure (without constraints) of the first 130 nucleotides of the RpfB 5’ UTR containing an intrinsic terminator structure. B: Northern blot of RNA from exponential and stationary phase cultures of *M. tuberculosis* and *M. bovis* BCG. RNA was separated by PAGE, transferred to a nylon membrane and probed with a ribo-probe indicated in Fig. 1B. C: Alternative structures of the RpfB 5’ UTR (1-130). The figure shows the two structures that were predicted with *mfold* (26) with the constraints that U117 is unpaired. Termination frequencies, (3’ RACE) have been indicated with bars. Point mutations that stabilise either the left conformation or the right conformation have been indicated with red and green circles, respectively. Red highlight indicates the terminator with the same sequence shown in green in its anti-terminated conformation.

To identify more precisely the 3’ termini associated with the RpfB 5’ UTR, we performed 3’ RACE as previously described (28). The results indicated that 12% of transcripts terminated well upstream of the poly-U tract, and 42% terminated within or proximal to the poly-U tract with U123 and U124 alone accounting for 17% (Fig. S2). Further downstream we found that 8% terminated at the newly annotated TTG, indicating transcriptional pausing associated with translation initiation, as recently reported for the TPP riboswitch (29).

The fact that more than a third of all 3’ termini fall within the poly-U tract, strongly favours the presence of a functional intrinsic terminator. The results also suggest that U117 is part of the poly-U tract and not the preceding stem as the structure in Fig. 3A suggests. We therefore re-modelled the RNA with the constraints that residues downstream of A116 were unpaired. This resulted in two alternatives; one with a slightly modified terminator structure and lower free energy than the original (ΔG -50.2 vs -49.6 kcal/mol); the other without terminator and with a higher free energy (ΔG -44.7 kcal/mol); we consider the latter a potential anti-terminated or read-through conformation (Fig. 3C).

In summary, our results indicate that the 5’ UTR of RpfB can adopt two conformations, one of which contains an intrinsic terminator, suggesting that this element comprises a novel RNA switch.

### Translational reporter fusions support the notion of an RNA switch

In order to verify and further characterise this putative, novel RNA switch we employed the previously described translational *lacZ* fusion. First, we compared constructs with and without the RNA switch, deleting the entire region from TSS1 to the end of the poly-U tract. This resulted in a significant increase in β-gal activity, suggesting that the RNA switch provides an additional layer of control by reducing RpfB expression during exponential growth (Fig. 4). The two conformations of the RpfB 5’ UTR are likely to exist in an equilibrium *in vivo.* We used these structures to predict single-nucleotide substitutions that could stabilise either conformation. Thus, a U6C substitution (green circles, Fig. 3C) would favour the anti-terminated structure, while a G112C substitution (red circles, Fig. 3C) would favour the terminated structure. The mutations were introduced into the *lacZ*-fusions and β-gal activity determined. The results (Fig. 4) demonstrate that stabilising the predicted terminator leads to significantly reduced *lacZ* expression (Fig. 4), while stabilising the antiterminator structure leads to significantly increased expression (Fig. 4). To further probe the intrinsic terminator, we made a mutant in which U117 to U119 were changed to adenines. This resulted in increased expression similar to that observed for the U6C mutant. These results substantiate the presence of the two structures and the potential to switch between these.

**Fig. 4:**
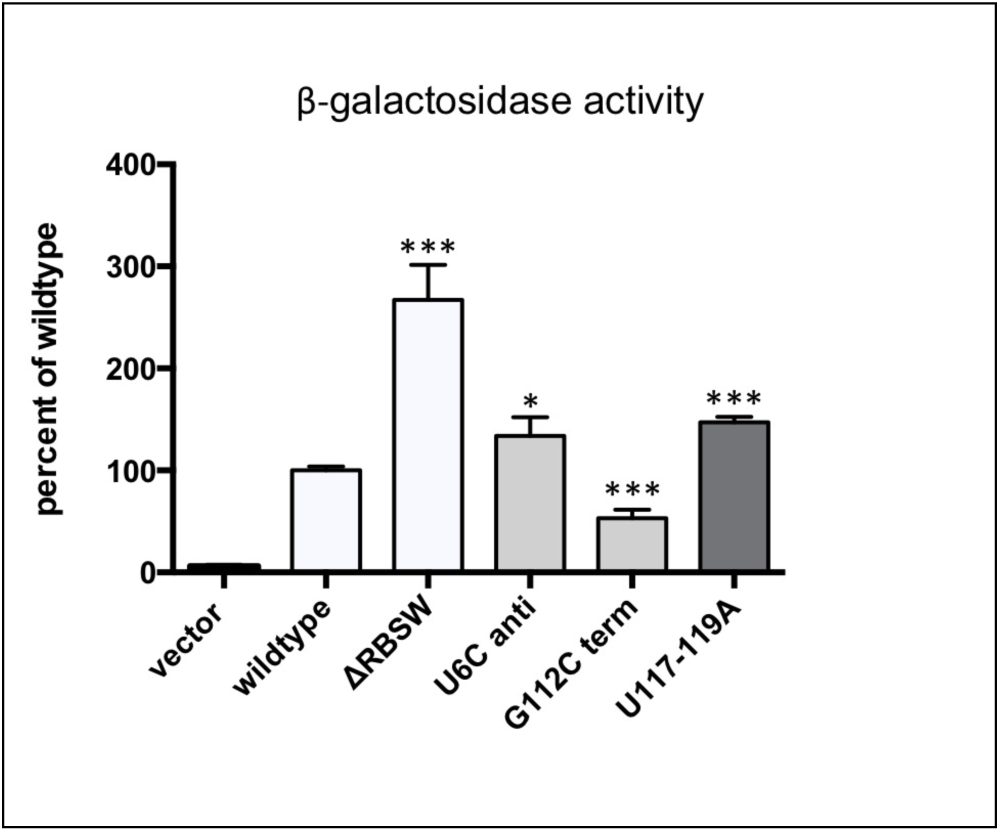
Reporter gene assays support the presence of an RNA switch. The figure shows β-galactosidase activity of translational reporters expressed in *M. smegmatis.* The constructs include the promoter region, 5’ UTR and 14 codons of the RpfB ORF, including ATG, as shown in Fig. 1. ΔRBSW: entire RNA switch deleted from the construct; U6C anti: point mutation predicted to stabilise the anti-terminated conformation; G112C term: point mutation predicted to stabilise the terminated conformation; U117-119A: change of three U residues to A residues. The values represent the mean and standard deviation of 6 biological replicates; *P ≤ 0.05; ***P ≤ 0.001.

Together our results strongly support that the RpfB 5’ UTR comprises a novel, transcriptional RNA switch that provides an additional layer of regulation to RpfB expression. As the terminated conformation has the lowest predicted free energy of the two, we assume it is the default conformation, and that its cognate ligand would promote read-through. Moreover, we find evidence of pausing associated with the start codon, which may provide even further control of RpfB expression.

### *rpfB, ksgA* and *ispE* form a tri-cistronic operon

Immediately downstream of the *rpfB* gene lies a gene encoding the highly conserved methyl transferase, KsgA that specifically methylates two adjacent adenosine residues in the 3’ end of the 16S ribosomal RNA (residues 1511 and 1512 within the sequence GGAAG in *M. tuberculosis*). This process is regarded as a checkpoint for ribosome maturation (30). There are no TSS identified between the *rpfB* 5’ UTR and the *ksgA* gene (Fig. 5), indicating that the two genes are part of the same operon. Moreover, according to the annotation, the ORFs for these two genes overlap, suggesting a very tight coupling in their expression. We tested if the two genes were co-transcribed using RT-PCR. The results, shown in Fig. 5 suggest that *rpfB* and *ksgA* are co-transcribed in both *M. tuberculosis, M. bovis* BCG and in the more distantly related *M. smegmatis.* In *M. tuberculosis,* but not in *M. smegmatis,* lies the the essential *ispE* downstream of *ksgA.* This gene encodes an essential ATP-dependent kinase involved in isoprenoid synthesis and ultimately, cell wall synthesis by providing the linker unit between arabinogalactan and peptidoglycan (31). Although there is a weak TSS 37 basepairs upstream of the annotated IspE GTG start codon as well as a consensus -10 motif (TAGTCT), we tested the possibility that *ispE* was co-transcribed with *rpfB* and *ksgA* due to the close proximity of the ORFs. The result, shown in Fig. 5, indicates that this is indeed the case and hence that *rpfB, ksgA* and *ispE* form a tri-cistronic operon in *M. tuberculosis* with an internal promoter driving baseline expression of *ispE.*

**Fig. 5:**
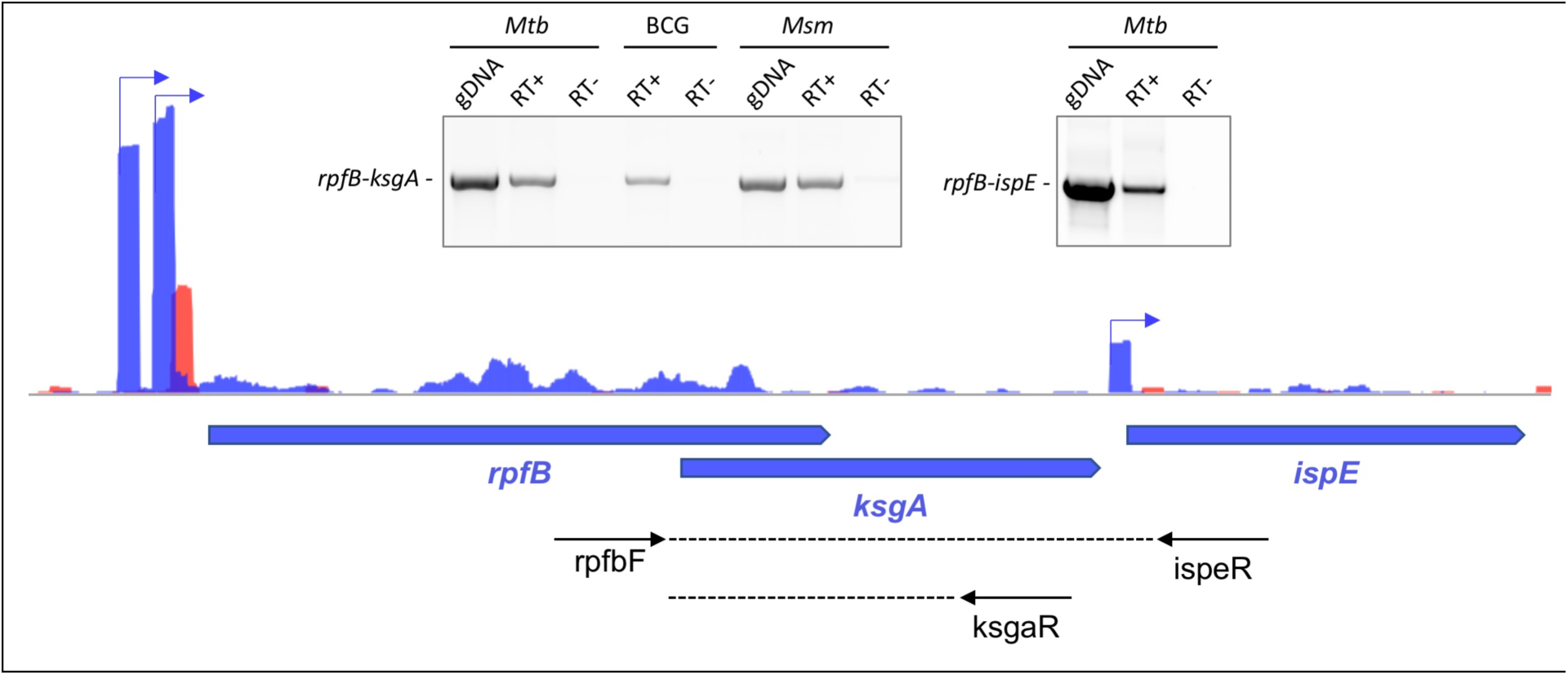
Co-transcription of *rpfB* with downstream genes. Main image shows three TSS associated with the *M. tuberculosis rpfB* locus on the plus strand; two for *rpfB* and a minor for *ispE,* according to global TSS mapping (22). Black arrows below locus indicate primers used for RT-PCR. Inserts show RT-PCR; left: *rpfB* and *ksgA* are co-transcribed in *M. tuberculosis* (*Mtb*), *M. bovis* BCG (BCG) and *M. smegmatis* (*Msm*); right: *rpfB*, *ksgA* and *ispR* are all co-transcribed in *Mtb*.

This, in turn indicates that rpfB, *ksgA* and *ispE* expression is regulated by the same RNA switch in *M. tuberculosis.* This arrangement provides a regulatory link between resuscitation, cell wall synthesis and ribosome maturation. It also offers the possibility that a cognate ligand could be associated with KsgA or IspE as well as with RpfB.

### Expression of RpfB during re-growth and nutrient starvation

To obtain a more detailed picture of termination and read-through of the RNA switch, we investigated the expression under different growth conditions. Initially we looked at expression as cells emerge from stationary phase into log-phase. A stationary phase culture, in which RpfB is poorly expressed, was diluted into fresh medium followed by RNA sampling over time. Fig. 6A shows a Northern blot of the time course probed for the RNA switch, which indicates robust expression of the terminated transcript after one hour in fresh medium, while expression of the longer, read-through transcripts reached a maximum later (around 5 hours) into the time course, suggesting that the cells require more time to achieve ligand concentrations permissive of read-through. We also investigated expression after the cells had been shifted to starvation conditions. Exponential phase culture was resuspended in nutrient deficient PBS+Tween80 followed by RNA sampling over time. Fig. 6B shows that P1-driven expression of RpfB ceases relatively quickly following nutrient starvation.

**Fig. 6:**
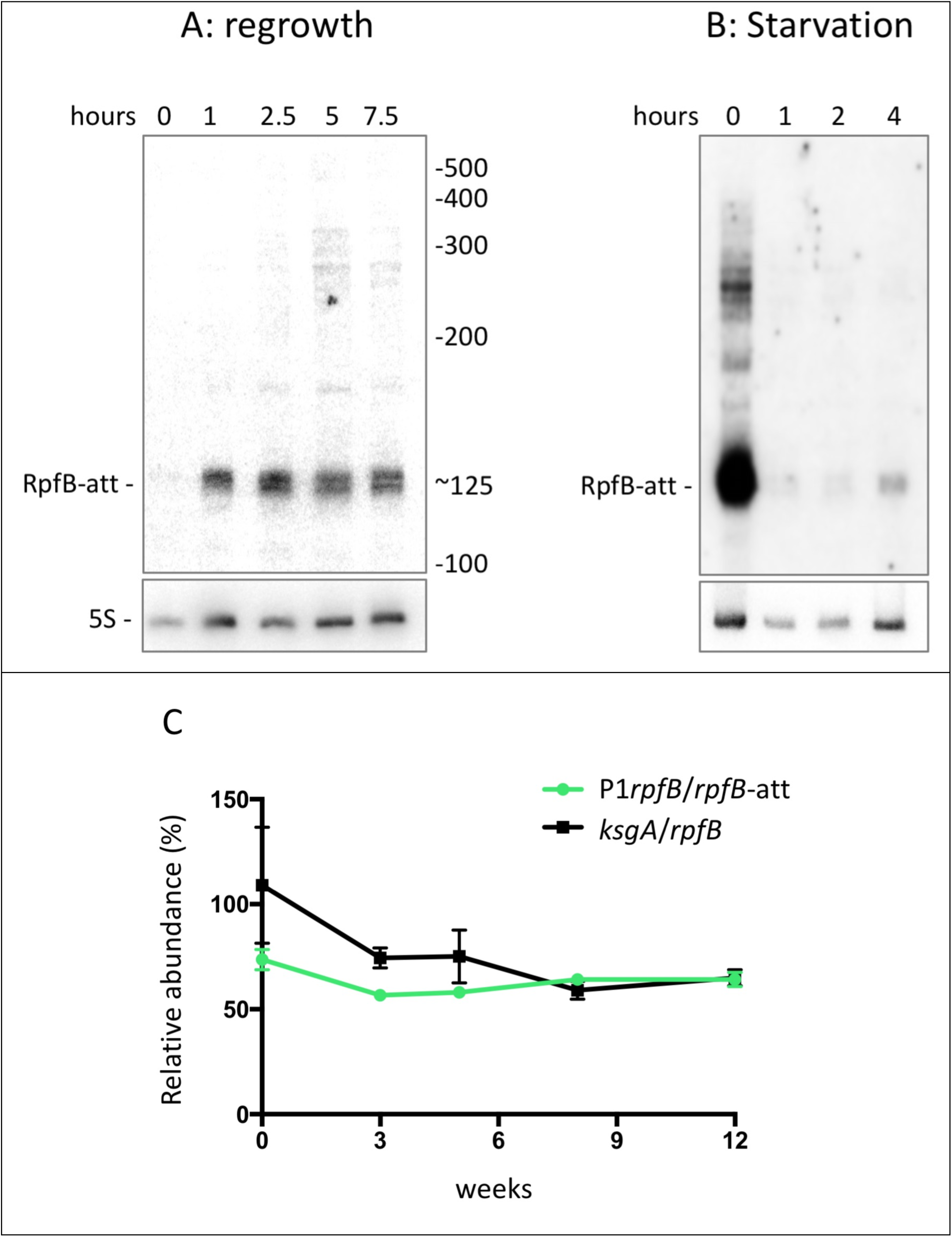
Expression and turnover of the RpfB 5’ UTR. Northern blots of *M. tuberculosis* RNA harvested at the indicated time points; RpfB-att corresponds to terminated transcript. A: after dilution of a stationary phase culture (1 week after OD_600_=1) into fresh medium. B: shows that expression of RNA switch ceases quickly after cells have been shifted to PBS + 0.05% Tween. For both 15 μg of RNA was separated by PAGE, transferred to a nylon membrane and probed for the RNA switch (oligos 1.48 for 5S RNA and 5.22 for RNA switch). C: Expression during biofilm formation. Normalised expression of P1 read-through and *ksgA* transcripts. Values for P1 derived *rpfB* mRNA (Fig. S4) were normalised to values for 5’ UTR RNA (P1rpfB/*rpfB-att),* and values for *ksgA* coding RNA were normalised to *rpfB* coding RNA. The graph illustrates the amount of P1-derived *rpfB* transcript relative to the amount of 5’ UTR transcript over 12 weeks of biofilm formation. Values represent mean and SD of three biological replicates.

### Expression of RpfB in biofilms

The formation of mycobacterial biofilms requires significant changes in gene expression followed by substantial re-arrangements of the cell wall (32); however, changes in Rpf expression have not been reported. We investigated the expression of the RNA switch as well as RpfB, asRpfB and KsgA in biofilms of *M. bovis* BCG, a close, more tractable relative of *M. tuberculosis* in which the entire *rpfB-ispE* transcript, including 5’ UTR, is 100% conserved. Biofilms were allowed to form in static, non-aerated cultures for the indicated period of time after which the pellicle was removed and processed for RNA. Quantitative real-time PCR (qRT-PCR) was performed for the 5’ UTR, P1 read-through, RpfB, asRNA and KsgA; details of these amplicons are outlined in Fig. S3, and the results are shown in Fig. S4. The results indicated that the level of all measured RNAs was slightly, but not significantly reduced during the initial stages of biofilm formation, but recovering as the biofilm matured. As *ispE* expression is driven by an additional promoter, we did not include this in our investigation.

In order to obtain values for transcriptional read-through vs termination within P1 derived transcripts, we normalised the raw values as outlined in Fig. S4. The final result, shown in Fig. 6C, indicates the level of P1-derived *rpfB* transcripts normalised to the values obtained for the 5’ UTR. The results can therefore be used as an approximation of the proportion of transcripts that proceed through the terminator region into the *rpfB* coding region. Similarly, we normalised the values for *ksgA* transcripts to *rpfB* transcripts to obtain a measure of relative abundance of the two cistrons. Overall the results indicate that there are no significant changes in the relative amounts of the investigated transcripts during biofilm formation.

### Transcription of the RpfB attenuator *in vitro*

In order to screen putative ligands of the RpfB RNA switch, we designed a single-round *in vitro* transcription assay. Since all four nucleotides are present within the first six positions of the RNA switch, we modified the 5’ end marginally to obtain a template that was suitable for single-round *in vitro* transcription (see Supplementary methods). We first tested the wildtype RNA switch and the three mutants from the reporter constructs, expressed from a heterologous promoter. Transcription read-through was observed either as template run-off or read-through SynB synthetic terminator (33). Halted elongation complexes were formed using *Escherichia coli* RNA polymerase (RNAP) and chased in the presence of heparin. The results demonstrated that the RpfB terminator is recognised by the *E. coli* RNAP resulting in approximately half of the complexes pausing/terminating at the predicted site (Fig. 7A, lanes 1 and 5), while the remaining continue transcription to obtain either the run-off transcript (lane 1) or the SynB terminated transcript (lane 5). Stabilising the terminator stem led to multiple signals around the RpfB terminator (lanes 2 and 6), while the run-off and the SynB terminated transcript were both replaced by aberrant signals that were approximately 30-40 nucleotides longer.

**Fig. 7:**
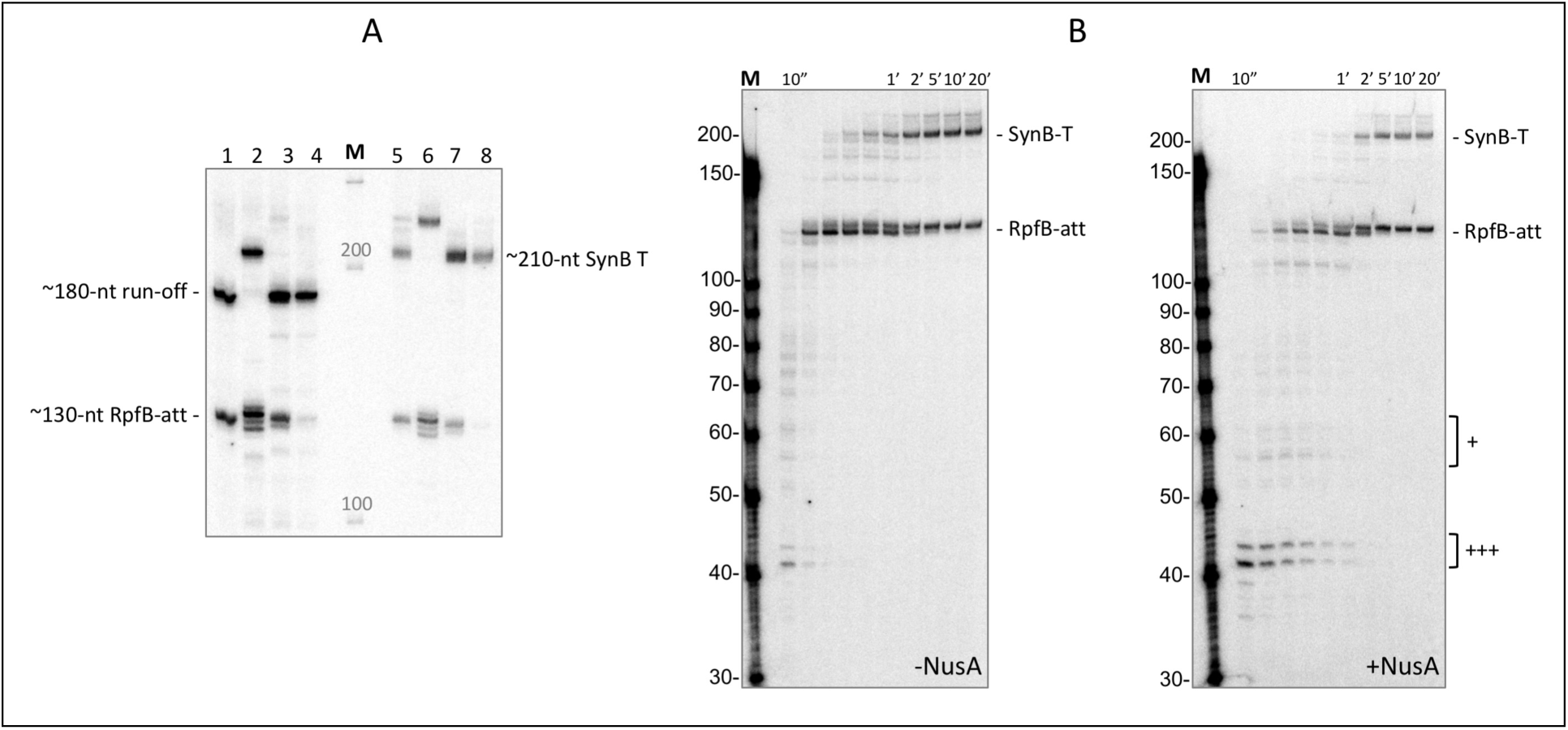
*In vitro* transcription of RNA switch. Transcription was initiated with GpU, omitting UTP from the initial reaction and labelling with 32P-aATP. Lanes 1-4 show reactions with run-off template; lanes 5-8 shows reaction from template with SynB-mediated termination instead of run-off. Lanes, 1+5: wildtype; 2+6: G112C term; 3+7: U6C antiterm; 4+8: U117-119A poly(A). B: Single-round *in vitro* transcription of the RpfB RNA switch. The RpfB RNA switch was transcribed *in vitro* with *E. coli* RNAP. Initiation complexes were stalled at position 11 and elongated in the presence of heparin and 50 μM NTP at 30°C. Left image is with RNAP only and right gel image is in the presence of 5-fold molar excess NusA. + symbols indicate regions with NusA enhanced pausing.

As this size transcript exceeded the theoretical maximum length possible using the template, we treated the samples with DNase to investigate the possibility of template labelling activity (not shown). However, this did not remove the aberrant signal, a phenomenon that we are currently unable to explain. More importantly, the two mutants, U6C and U117-119A both displayed decreased termination at the RpfB terminator and increased read-through, supporting the *in vivo* findings and lending significant support to the presence of a transcriptionally regulated RNA attenuator (Fig. 7A, lanes 3, 4, 7 and 8). Some transcriptionally regulated RNA attenuators require the RNAP to pause at specific sites to allow co-transcriptional folding and ligand binding (34). We investigated the pausing pattern of the RNA switch in a time-course experiment in the presence and absence of NusA, a transcription factor known to promote transcriptional pausing. As suspected, there were several pause sites within the sequence, most of which were enhanced in the presence of NusA, resulting in an overall reduced elongation rate (Fig 7B). We observed a particularly enhanced pause signal around position 41 and 43, corresponding to positions 37 and 39 in the true RNA switch transcript (indicated with +++ in Fig. 7B).

These results suggest that *in vivo,* NusA may be required to allow more time for potential ligand interactions which may be necessary for anti-terminator formation and transcriptional read-through.

### The RpfB attenuator is restricted to a subset of pathogenic mycobacteria

Many riboswitches are highly conserved, particularly between closely related species. To investigate the occurrence of the newly identified RpfB attenuator, we aligned sequences upstream of the RpfB coding region from seven mycobacterial species (Fig. 8). The alignment indicated that the P2 -10 region is identical in all of the selected species.

**Fig. 8:**
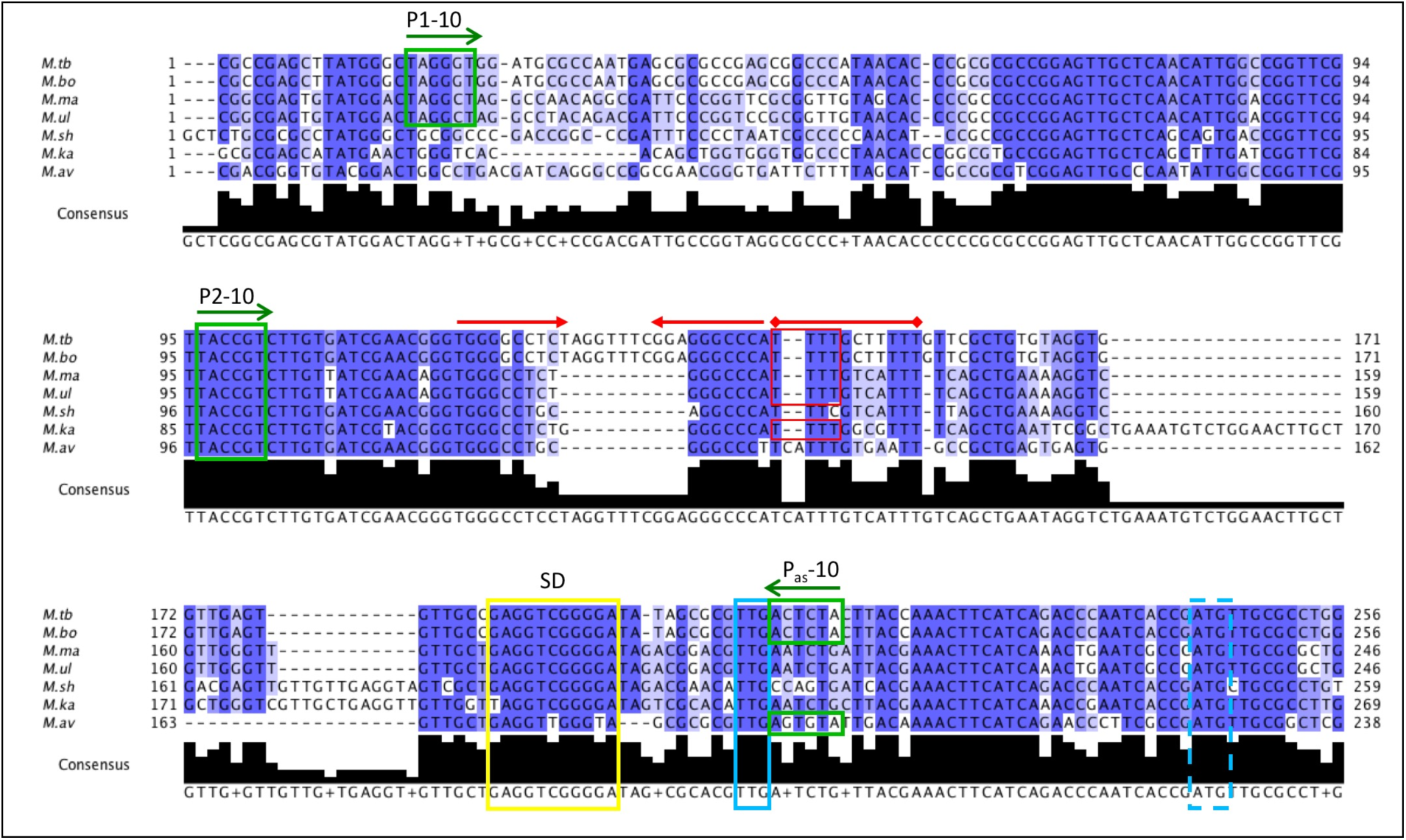
Sequence alignment of *rpfB* promoter regions and 5’ UTRs. Selected *Mycobacterium spp* are aligned: *M. tuberculosis* (*M.tb*), *M. bovis* (*M.bo*), *M. marinum* (M.ma), *M. ulcerans* (*M.ul*), *M. shinjukuense* (*M.sh*), *M. kansasii* (*M.ka*), *M. avium* (*M.av*); an extended alignment can be seen in Fig. S5. The alignment is coloured by % sequence identity (darker blue = higher conservation). Green boxes: -10 regions; blue box: TTG start codon; blue dashed box: previously annotated ATG start; yellow box: SD sequence. Red arrows: inverted repeats, followed by red line indicating poly-U tract based on *M. tb* sequence.

P1, on the other hand, is less well-conserved, and the TANNNT -10 consensus indicative of a functional promoter is only seen in *M. tuberculosis, M. bovis, M. marinum and M. ulcerans,* suggesting that the long 5’ UTR is restricted to a subset of pathogenic mycobacteria; additional species from the MTBC, including *Mycobacterium africanum, Mycobacterium microti* and *Mycobacterium canetti* had sequences that were identical to *M. tuberculosis* (not shown). A more extensive alignment is shown in Fig. S5. To investigate the termination potential within the selected species, regardless of P1, we analysed the region using Transterm (35) to identify putative hairpins/terminators. Only *M. tuberculosis, M. bovis, M. marinum, M. ulcerans* and *M. kansasii* have hairpins followed by at least four consecutive uracil residues, indicative of a functional terminator (33,36) (Fig. S6). Thus, on the basis of promoter and terminator motifs we conclude that this RNA attenuator is only present in a subset of pathogenic mycobacteria with a phylogenetic split between MBTC/*M. marinum/M. ulcerans* and *M. leprae,* which is consistent with the split seen by aligning 16S sequences (37). Moreover, only *M. tuberculosis, M. bovis* and *M. avium* appear to have a functional antisense promoter, indicating that the presumably tight regulation provided by multiple promoters, RNA attenuator and asRNA is specific for species within the MTBC. The association of this element with certain pathogenic species only, offers the possibility that its function is associated with pathogenesis and adaptation to the host environment.

## Discussion

Rpfs are cell wall remodelling enzymes with the potential to lyse and kill the cells that express them. Hence, their expression is under tight often multi-layered control, and we propose disruption of such control as a target for drug development. In the current study, we have shown that multipronged regulation also applies to the expression of *M. tuberculosis* RpfB. This gene is transcribed from two promoters and post-transcriptionally regulated by an entirely novel pathogen-specific, transcriptionally regulated RNA switch. Moreover, we provide evidence for a functional antisense promoter, which may regulate RpfB expression by RNA polymerase collision, inhibition of translation initiation, mRNA processing or all of the above. We also show that this level of multi-layered control appears to be specific for species within the MTBC, as one or more of the described elements are absent from other mycobacterial species. Moreover, we show that *rpfB, ksgA* and *ispE* form a tri-cistronic operon, implying that all of these regulatory mechanisms may extend to *ksgA* and *ispE* expression as well. However, *ispE* is essential and expression likely to be affected in the previously described *rpfB* deletion strain (38). Hence, we assume that the weak TSS upstream of *ispE* is sufficient for survival, while the tri-cistronic arrangement with *rpfB* and *ksgA* ensures coordinated expression of the genes directed by the RNA switch. In addition, we have re-annotated the translation start site and identified a likely ribosome-binding site based on several lines of experimental evidence. Finally, we also found evidence of 'start codon associated pausing’, which has recently been shown to have importance for riboswitch-regulated gene expression (29).

In contrast to the relatively well-conserved genetic arrangement of *tatD-rpfB-ksgA*, the RNA switch is restricted to a small subset of pathogenic mycobacteria, including *M. tuberculosis.* Based on predictions of structure and free energy, we expect that a cognate ligand increases transcriptional read-through.

Our *in vitro* transcription assays demonstrate that there are several pause sites within the RNA switch region and that these are enhanced by NusA. We expect that these pauses may be critical for co-transcriptional folding and ligand recognition.

Some of our results did not agree with those previously published (25). We did investigate the possibility of an additional promoter downstream of P2, but our reporter gene fusions and previously published dRNA-seq (22) confirm that there is no promoter activity in that region. We employed two different means of determining the translation start and findings were further supported by the presence of a likely SD sequence. Therefore, we regard TTG as the correct start site. This also means that the asRNA is positioned immediately upstream of the start codon and covering the newly identified SD sequence.

At present, we have not identified a ligand for this RNA switch, although several have been tested in our *in vitro* transcription assay. The different growth conditions tested here do not allude to any specific ligand. It remains a possibility that the switch between termination and anti-termination is not mediated by a small molecule, characteristic of a *bona fide* riboswitch, but rather a protein ligand, similar to ribosomal protein operons or yet another molecule/mechanism capable of pushing the equilibrium between the two conformations. The lack of widespread conservation and the fact that the genes regulated by this element are not associated with metabolic pathways further complicates the prediction of this ligand. The coordinated expression of RpfB, KsgA and IspE fits a model in which one or more molecular signals leading to resuscitation and cell wall remodelling/synthesis associated with growth, also lead to activation of protein synthesis by allowing the final steps in ribosome maturation. Moreover, the coordinated expression of the genes within this operon ensures that the cell maintains a carefully balanced ratio between different aspects of macromolecular synthesis, which is also apparent in operons encoding RNA polymerase subunits together with ribosomal proteins. The regulatory link described in this study means that resuscitation and ribosome maturation or rephrased, cell wall synthesis and protein synthesis, two classical antimicrobial targets could be simultaneously targeted via the RpfB RNA switch. Disruption of the regulation of its action provides an opportunity for development of a novel class of anti-tubercular drugs with a unique mode of action.

## Materials and methods

### Bacterial strains and growth conditions

*E. coli* DH5α were grown in LB liquid media or agar (1.5%) supplemented with 250 μg/mL hygromycin B as required.

*Mycobacterium smegmatis* mc^2^155(39) was grown on LB agar supplemented with 50 μg/mL hygromycin B as required, and in liquid LB media supplemented with 0.05% Tween 80 and 50 μg/mL hygromycin B as required. *M. tuberculosis* H37Rv (40) and *M. bovis* BCG were grown on Middlebrook 7H11 agar supplemented with 10% OADC, 0.5% glycerol and 50 μg/mL hygromycin B as required and in liquid Middlebrook 7H9 medium supplemented with 10% ADC, 0.4% glycerol and 0.05% Tween 80 in roller bottles (Cell Master, Griener Bio-One) or PETG flasks (Nalgene, Thermo Scientific), respectively. Exponential phase cultures were harvested at OD_600_ 0.6-0.8 Stationary phase cultures for *M. tuberculosis* and *M. bovis BCG* were harvested at least 1 week after 1.0 OD_600_. For time-course experiments, cultures were harvested as indicated. Biofilms were formed by adding 10 ml of an exponential phase culture to 50 ml polypropylene tubes, sealing tightly and leaving for the indicated amount of time. At time of harvest the pellicle was removed and processed for RNA. Mycobacteria were transformed by electroporation.

### Plasmid construction

Plasmids used in this study are listed in Table S1.

### Oligonucleotides

Oligonucleotides used during this study are listed in Table S2.

### RNA isolation

Total RNA extraction was performed as previously described(28).

### cDNA synthesis and 3’ rapid amplification of cDNA ends (RACE)

cDNA was synthesised using Superscript III reverse transcriptase (Invitrogen), largely according to manufacturer protocol except for an additional extension step for 30 minutes at 55°C, priming reactions with random hexamers (Promega).

3’ RACE was performed as previously described (41). Samples were reverse transcribed and primed using oligo d(T) adapter primer (oligonucleotide 2.07). RACE targets were amplified using adapter primer and a gene specific primer (oligonucleotide 2.09 and 2.15 respectively).

Co-transcription of *rpfB* and *ksgA* was analysed using cDNA generated with random hexamers. cDNA was amplified with REDTaq PCR reaction using primers flanking the *rpfB/ksgA* overlapping region complementary to the sequence in *M. tuberculosis, bovis* BCG and *smegmatis* (oligonucleotides 5.51 and 8.04).

### Northern blotting

Northern blotting and probing was performed as described in (41).

Template oligonucleotides are listed in Table S2. Membranes were exposed to a phosphor screen and developed using Typhoon FLA 9500 (GE), sizing RNAs using Century marker (Ambion).

### Quantitative RT-PCR

‘SensiFast SYBR Hi-ROX master mix’ (Bio-line) was used to amplify cDNA for quantitative RT-PCR (qRT-PCR), according to manufacturer’s instructions. *M. tuberculosis* H37Rv DNA was used to create a standard curve. Wells were loaded with either 1 μL standard in three technical replicates or 1 μL RT+/- cDNA in 4 technical replicates.

All reactions were carried out using a ‘QuantStudio 6 Flex Real-Time PCR System’ and analysed using QuantiStudio Real-time PCR software v1.1 (Applied Biosciences).

### β-galactosidase assay

Protein extracts were obtained from cultures of *M. smegmatis* and assayed as previously described(28).

Miller units were expressed as a percentage of the average WT value. Statistical significance was calculated using one-way ANOVA with Tukey post hoc analysis in IBM SPSS ±1 standard deviation. Significance thresholds specified as: ‘NS’ (no significant difference, P > 0.05), ‘*’ (P ≤ 0.05), ‘**’ (P ≤ 0.01) and ‘***’ (P ≤ 0.001).

### *In vitro* transcription

*E. coli* RNAP *in vitro* transcription assays were carried out using previously described methods for producing halted transcription elongation complexes (TECs)(42,43). Transcription templates were cloned into pGAMrnnX (see supplemental methods for details).

### Q5 site directed mutagenesis

Site-directed mutagenesis (SDM) was carried out using the ‘Q5 SDM kit’ (NEB) following the manufacturer protocol. Correct constructs were sub-cloned into un-treated vector.

### Overlap extension mutagenesis

For small mutations, a pair of Phusion GC polymerase PCR reactions were carried out: reaction (A) used an upstream forward primer and a mutagenic reverse primer spanning the region to be mutated, reaction (B) used a downstream reverse primer and a mutagenic forward primer spanning the region to be mutated. The resulting amplicons contained a region of complementarity exploited in reaction (C) by combining 1 μL of each as template in another PCR reaction with the non-mutagenic primers of the original reactions.

### Alignment of the *tatD-rpfB* intergenic regions

Test alignments of the intergenic regions between the *tatD* and *rpfB* genes in a number of mycobacteria indicated that a small number of mycobacterial species aligned well whereas others had large insertions and deletions. Based on the preliminary alignments, we selected a number of species with well-conserved intergenic regions to align first (Fig. 7) and subsequently added to this alignment three more species to highlight divergence in the sequences of the latter (Fig. S6). Further details can be found in supplementary methods.

## Acknowledgements

We thank Galina Mukamolova and Jeff Green for helpful discussions and Finn Werner for critical reading of the manuscript.

